# Kingdom-wide comparison reveals conserved diurnal gene expression in Archaeplastida

**DOI:** 10.1101/387316

**Authors:** Camilla Ferrari, Sebastian Proost, Marcin Janowski, Jörg Becker, Zoran Nikoloski, Debashish Bhattacharya, Dana Price, Takayuki Tohge, Arren Bar-Even, Alisdair Fernie, Mark Stitt, Marek Mutwil

## Abstract

Plants have adapted to the diurnal light-dark cycle by establishing elaborate transcriptional programs that coordinate innumerable metabolic, physiological, and developmental responses to the external environment. These transcriptional programs have been studied in only a few species, and their function and conservation across algae and plants is currently unknown. We performed a comparative transcriptome analysis of the diurnal cycle of nine members of Archaeplastida, and we observed that, despite large phylogenetic distances and dramatic differences in morphology and lifestyle, diurnal transcriptional programs of these organisms are similar. However, the establishment of multicellularity coincided with the uncoupling of cell division from the diurnal cycle and decreased diurnal control of the expression of the biological pathways. Hence, our study provides evidence for the universality of diurnal gene expression and elucidates its evolutionary history among different photosynthetic eukaryotes.

## Introduction

The endosymbiotic event that occurred between a eukaryotic ancestor and cyanobacteria led to the formation of the plastid and emergence of early single-celled members of the Archaeplastida. These organisms established multicellularity, conquered land, and evolved complex organs to adapt and survive the new environmental challenges (Chang *et al.*, 2016). Comparative genomic studies aim to reveal how changes in genomes and gene families have facilitated the evolution of new features (Guo, 2013; Bowman *et al.*, 2007; Proost *et al.*, 2009; Goodstein *et al.*, 2012) and have led to important discoveries. These include the single origin of the plastid in eukaryotes (Price *et al.*, 2012) and the emergence and expansion of gene families that are essential for drought tolerance and for hormone synthesis during the appearance of land plants (Rensing *et al.*, 2008). However, on its own, genome sequence data might not reveal which genes work together towards establishing a trait, and how new traits are formed from new or existing genes (Ruprecht *et al.*, 2017a; Ruprecht *et al.*, 2017b; Ruprecht *et al.*, 2016). To address this paucity of functional information from genome data, researchers compare other gene functional data, such as protein-protein interactions (Vazquez *et al.*, 2003) and gene expression (transcriptomic) data (Wu *et al.*, 2002). Comparative transcriptomics were used to identify molecular mechanisms underlying interspecific differences related to plant adaptation (Nakayama *et al.*, 2018) and to identify the evolution of metabolic responses in microalgae (Zhou *et al.*, 2017). Furthermore, comparative transcriptomics can be used in a broader fashion to discover shared and unique events that occurred in evolution. This can be accomplished by comparing distantly related species, as exemplified by the identification of modules that are involved in development and that are shared across human, worm, and fly (Gerstein *et al.*, 2014) and by the discovery of conserved early and late phases of development in vertebrates (Levin *et al.*, 2016).

Photosynthetic organisms are capable of perceiving day length and the alternation of light and dark through two primary systems: light-dark detection and the circadian clock (Serrano-Bueno *et al.*, 2017). In addition, light acts as the energy source that drives photosynthesis. The circadian clock, light signalling, and the supply of energy and fixed carbon are the driving forces of metabolism and of many biological processes whose expression is regulated by alternating light and dark intervals (Dodd, 2005; B. Usadel *et al.*, 2008). Many studies have addressed how organisms perform under different diurnal conditions and how they coordinate the activity of their biological pathways. Diurnal regulation actively controls physiological processes, such as cell division, metabolic activity, and growth in all domains of life (Covington *et al.*, 2008; Stockel *et al.*, 2008; Bell-Pedersen *et al.*, 2005; Zones *et al.*, 2015; de los Reyes *et al.*, 2017). Gene expression plays a fundamental role in controlling the activity of biological pathways and is under strict diurnal regulation. For example, in *Ostreococcus tauri* most of the genes are under control of light/dark cycles and specifically, genes involved in cell division, the Krebs cycle and protein synthesis, have the highest amplitude within the diurnal cycle (Monnier *et al.*, 2010; de los Reyes *et al.*, 2017). Studies in *Drosophila melanogaster* show that in addition to gene expression, splicing, RNA editing and non-coding RNA expression are highly affected by diurnal regulation (Hughes *et al.*, 2012), whereas studies in mammals highlight the importance of diurnal regulation when applied to pharmaceutical applications (Zhang *et al.*, 2014; Archer *et al.*, 2014; Mure *et al.*, 2018). Furthermore, recent studies on plants have shown a moderate level of conservation of diurnal responses among distantly related species, along with occasional divergent evolution of specific mechanisms (de los Reyes *et al.*, 2017; Serrano-Bueno *et al.*, 2017).

In this study, we analysed diurnal transcriptomes of distantly related species of the Archaeplastida, including eukaryotic algae, different phylogenetic groups of terrestrial plants, and a cyanobacterium as an outgroup. Our overarching goal was to understand how diurnal gene expression has evolved to accommodate the appearance of new gene families, morphologies, and lifestyles. To this end, we generated, collected and compared diurnal gene expression data of model organisms representing nine major clades of photosynthetic eukaryotes, comprising *Cyanophora paradoxa* (glaucophyte, early diverging alga), *Porphyridium purpureum* (rhodophyte), *Chlamydomonas reinhardtii* (chlorophyte)*, Klebsormidium nitens* (charophyte), *Physcomitrella patens* (bryophyte), *Selaginella moellendorffii* (lycophyte), *Picea abies* (gymnosperm), *Oryza sativa* (monocot), and *Arabidopsis thaliana* (eudicot). We demonstrate that diurnal transcriptomes are significantly similar despite the large evolutionary distances, morphological complexity, and habitat. We find that whereas cell division of the single-cellular and simple multicellular algae is synchronized by light, this behaviour is lost in land plants. Establishment of multicellularity has also resulted in a less defined diurnal control of most biological processes. Finally, we show that the core components of the circadian clock show conserved expression among plant clades, thus suggesting a potential mechanism behind conserved diurnal gene expression. Our study represents a significant advance in our understanding of how diurnal gene expression evolved in the anciently derived Archaeplastida.

## Results and Discussion

### Identification of diurnally regulated genes in the Archaeplastida

To generate comprehensive diurnal expression atlases capturing the evolution of the Archaeplastida, we retrieved publicly available diurnal gene expression data from *Synechocystis* sp. PCC 6803 (Beck *et al.*, 2014), *Chlamydomonas reinhardtii* (Zones *et al.*, 2015), *Oryza sativa* (Xu *et al.*, 2011), and *Arabidopsis thaliana* (Blasing, 2005) and generated new RNA sequencing data from *Cyanophora paradoxa*, *Porphyridium purpureum*, *Klebsormidium nitens*, *Physcomitrella patens*, *Selaginella moellendorffii*, and *Picea abies* (Fig. 1A). To generate the data, we subjected *P. purpureum*, *K. nitens*, *P. patens*, *S. moellendorffii* to a 12-hour light / 12-hour dark photoperiod (12L/12D) and *C. paradoxa*, *P. patens*, and *P. abies* to a 16L/8D photoperiod. Different photoperiods where used because for species such as *P. abies* (herein referred to as spruce), a shorter photoperiod would cause dormancy and growth cessation (HEIDE, 1974; Singh *et al.*, 2017). The difference in photoperiod was shown to have a relatively small impact on the organization of daily rhythms (de los Reyes *et al.*, 2017). Starting from ZT1 (Zeitgeber time), defined as one hour after the onset of illumination, we extracted RNA from the six organisms every two hours, resulting in twelve time points per species. We obtained an average of 20 million reads per sample (Supplementary Table 1), whereby more than 85% of the reads mapped to the genome, and on average 70% of the reads mapped to the coding sequences. Principal Component Analysis (PCA) of the samples showed separation according to the time and between light and dark samples for most of the species, with exception of spruce (Supplementary Fig. 1).

**Fig. 1.**
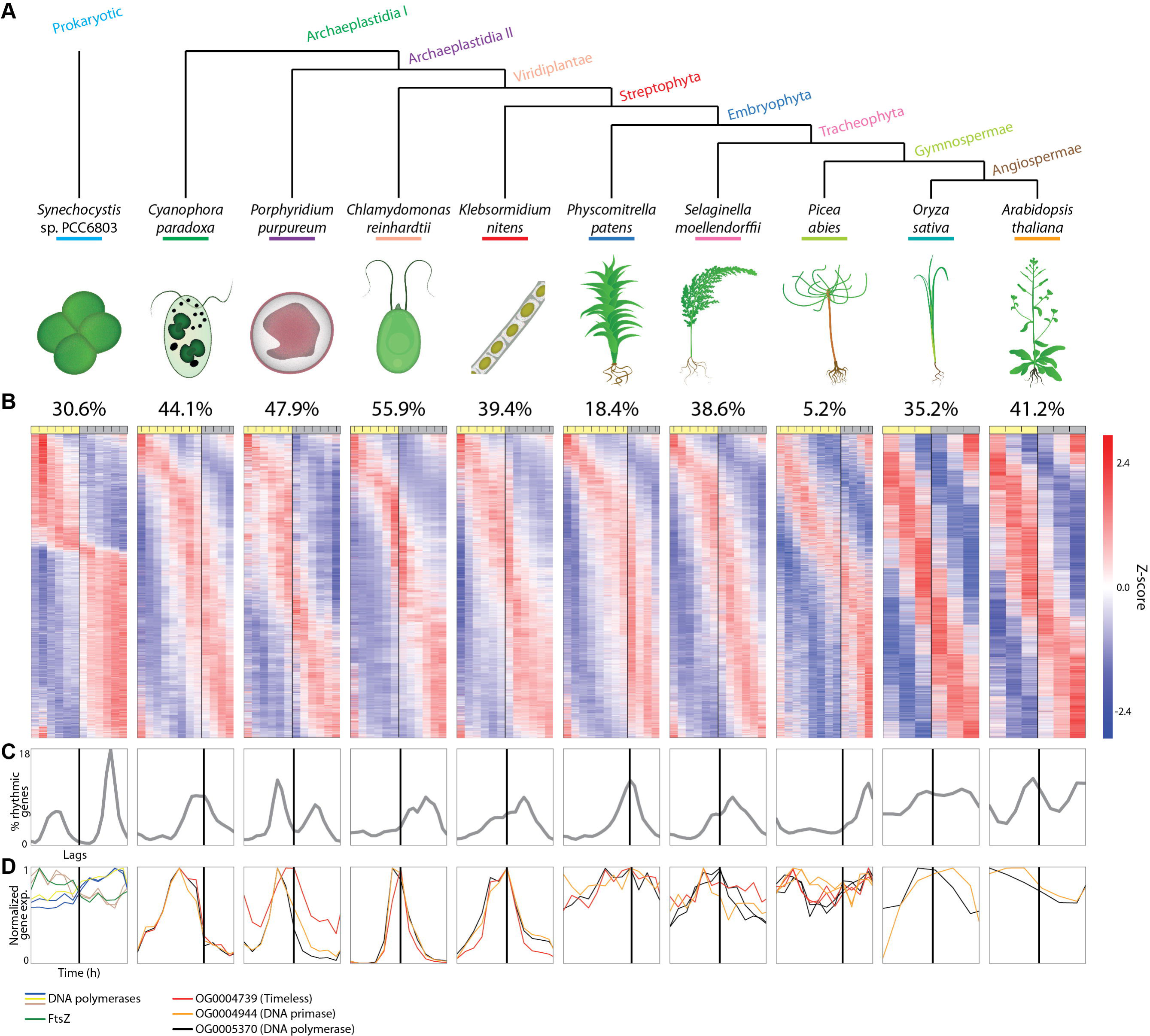
Diurnal gene expression of plant clades. A) Species tree of the Archaeplastida, together with the respective phylostratum of each clade. Synechocystis is included as a cyanobacterial outgroup. B) The percentage of rhythmic genes in a given species is shown above the yellow (light) and grey (darkness) bar indicating the photoperiod of each diurnal experiment. Below, z-score standardized gene expression profiles for each species sorted by the lag, signifying the time point of highest expression during the day. C) The distribution of lags of rhythmic genes through the diurnal experiment. D) Expression profiles of cell division genes. The coloured lines indicate the different gene families. Each family can have more than one gene.

To detect genes that are regulated by the diurnal cycle and to elucidate the time of the day where these genes reach peak expression, we set out to identify genes showing rhythmic gene expression. To this end, we used JTK_Cycle algorithm (Hughes *et al.*, 2012), which revealed significantly rhythmic genes and the ‘lag’ values of these genes, which ranges from 1 (the gene peaks at the first time-point of the series) to 24 (the gene peaks at the last time-point, Supplementary Table S2-11).

We observed that between 5.2% - 55.9% of genes show significantly rhythmic expression across species (adjusted *p*-value < 0.05, Fig. 1B). We observed that single-celled algae and multicellular land plants tend to be on average more rhythmic (on average 49.3% genes rhythmic for Cyanophora, Porphyridium, and Chlamydomonas), than multicellular land plants (on average 33.3% rhythmic genes for Physcomitrella, Selaginella, rice, and Arabidopsis). We excluded spruce from this calculation because this species showed an unusually low level of rhythmic gene expression (5.2%). However, this is in line with studies on diurnal expression of photosynthetic genes, which are largely rhythmic in flowering plants, but not in spruce and other gymnosperms (Brinker *et al.*, 2001; Gyllenstrand *et al.*, 2014; Oberschmidt *et al.*, 1995). This suggests that diurnal gene expression is not crucial in gymnosperms, presumably as a consequence of extreme variation in day length at high latitudes, and the long photoperiod during the growing season (Dormling *et al.*, 1968; HEIDE, 1974).

To elucidate when rhythmic genes peak during the day, we plotted the number of genes assigned to each lag (Fig. 1C). From glaucophyte to gymnosperm, the majority of rhythmic genes tend to peak during the second half of the day, at either dusk or night. The exception to this trend is the red alga Porphyridium, in which rhythmic genes appear in two peaks, in the middle of the light and dark period, similarly to the cyanobacterium *Synechocystis* sp. PCC 6803. Furthermore, in angiosperms, we observed a more uniform distribution of rhythmic genes with double peaks coinciding with transitions from day to night and from night to day (Fig. 1C). The more uniform expression suggests that, in contrast to algae and early-diverging land plants, gene expression in angiosperms is less under the control of the diurnal cycle.

The diurnal cycle is known to synchronize cell division in cyanobacteria (Vaulot *et al.*, 1995), red algae (Suzuki *et al.*, 1994), and green algae (Lien and Knutsen, 1979), and cell division is accompanied by a sharp increase in the transcript levels of DNA polymerases, primases, and other enzymes (Monnier *et al.*, 2010; Zones *et al.*, 2015). The expression of these marker genes revealed that single celled algae and the relatively simple multicellular Klebsormidium exhibit a single expression peak of cell division genes at the onset of darkness (Fig. 1D). Conversely, the more complex multicellular land plants show ubiquitous expression of these genes (Fig. 1D). This suggests that the increased morphological complexity of multicellular land plants necessitated uncoupling of the diurnal cycle and cell division.

### Phylostrata analysis of rhythmicity and expression

The age of appearance of gene families is positively correlated with their expression levels and severity of the corresponding mutant phenotypes (C. Ruprecht *et al.*, 2017). To investigate whether there is a relationship between the age of a gene family and its diurnal expression pattern, we first defined the age of gene families with a phylostratigraphic analysis, which identifies the earliest clade found within a gene family (Guo, 2013). Because the order of appearance of the clades is known (Fig. 1A), gene families can be sorted according to their relative age (phylostrata), from oldest phylostratum (i.e., present in cyanobacteria and all Archaeplastida) to youngest (e.g., specific to only Arabidopsis or rice, Supplementary Table 12). This analysis revealed that the majority of genes are old and belong to the earliest phylostratum for algae and plants (Fig. 2A, green bar). Next, we plotted the distribution of expression peaks of the rhythmic genes from the different phylostrata, and observed similar diurnal expression patterns for all phylostrata (Fig. 2B, an example from *P. patens*, Supplementary Fig. 2 for other species), indicating that the age of genes does not strongly influence the diurnal timing of gene expression.

**Fig. 2.**
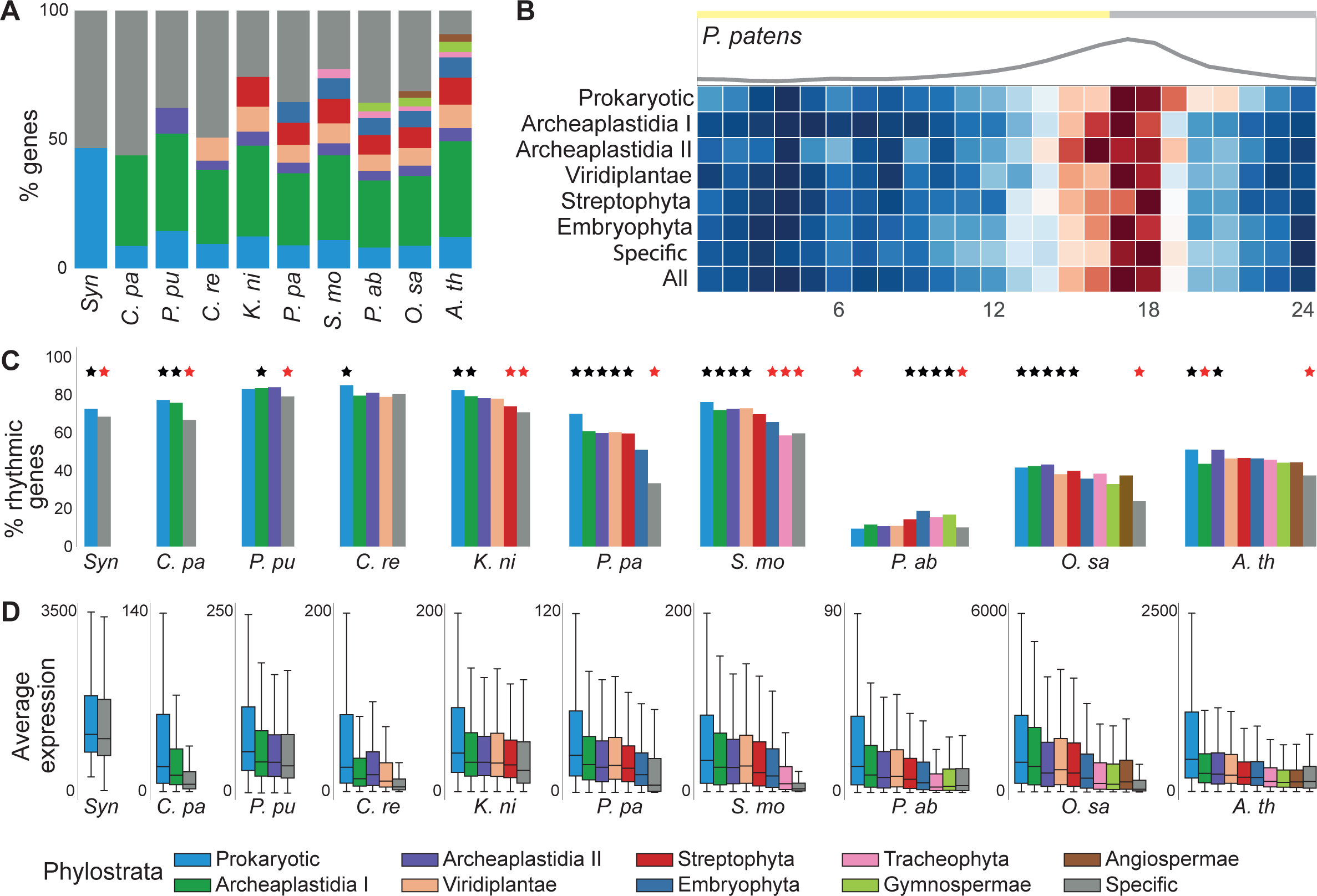
Expression analysis of phylostrata. A) Percentages of genes in each phylostrata per species. B) Distribution of lags for the phylostrata in Physcomitrella. The plot above the heat map shows the cumulative distribution of the lags. C) Analysis of phylostrata rhythmicity per species. The bars represent the percentage of rhythmic genes for each phylostratum. Black and red stars indicate phylostrata that are significantly enriched or depleted for rhythmic genes, respectively (FDR corrected empirical p-value < 0.05). D) Average expression of genes assigned to the phylostrata in the different species.

We also explored if there is a relationship between the age of gene families and their rhythmicity, by comparing the percentage of significantly rhythmic genes within each phylostratum. There was a significant tendency for genes from older phylostrata to have higher rhythmicity than genes from younger phylostrata (Fig. 2C). For example, the oldest phylostratum, Prokaryotic (containing genes found in bacteria and eukaryotes, light blue boxplot), was more rhythmic than expected by chance for most of the analysed organisms (Prokaryotic and Archaeplastida I phylostrata, FDR corrected empirical *p*-value < 0.05). Conversely, the youngest phylostrata, comprising species-specific or clade-specific genes (Specific phylostratum, grey bar, FDR corrected empirical *p*-value < 0.05) were less rhythmic than expected by chance for all species, with the exception of Chlamydomonas and spruce.

Finally, we asked whether the age of a gene family influences the expression levels of the transcripts, and observed that the gene expression levels significantly decrease as the phylostrata become younger, and the lowest expression is reported for species-specific or clade-specific gene families (Fig. 2D, FDR corrected *p*-value < 0.05, Supplementary Fig. 3). We therefore conclude that older phylostrata tend to be more strongly expressed.

**Fig. 3.**
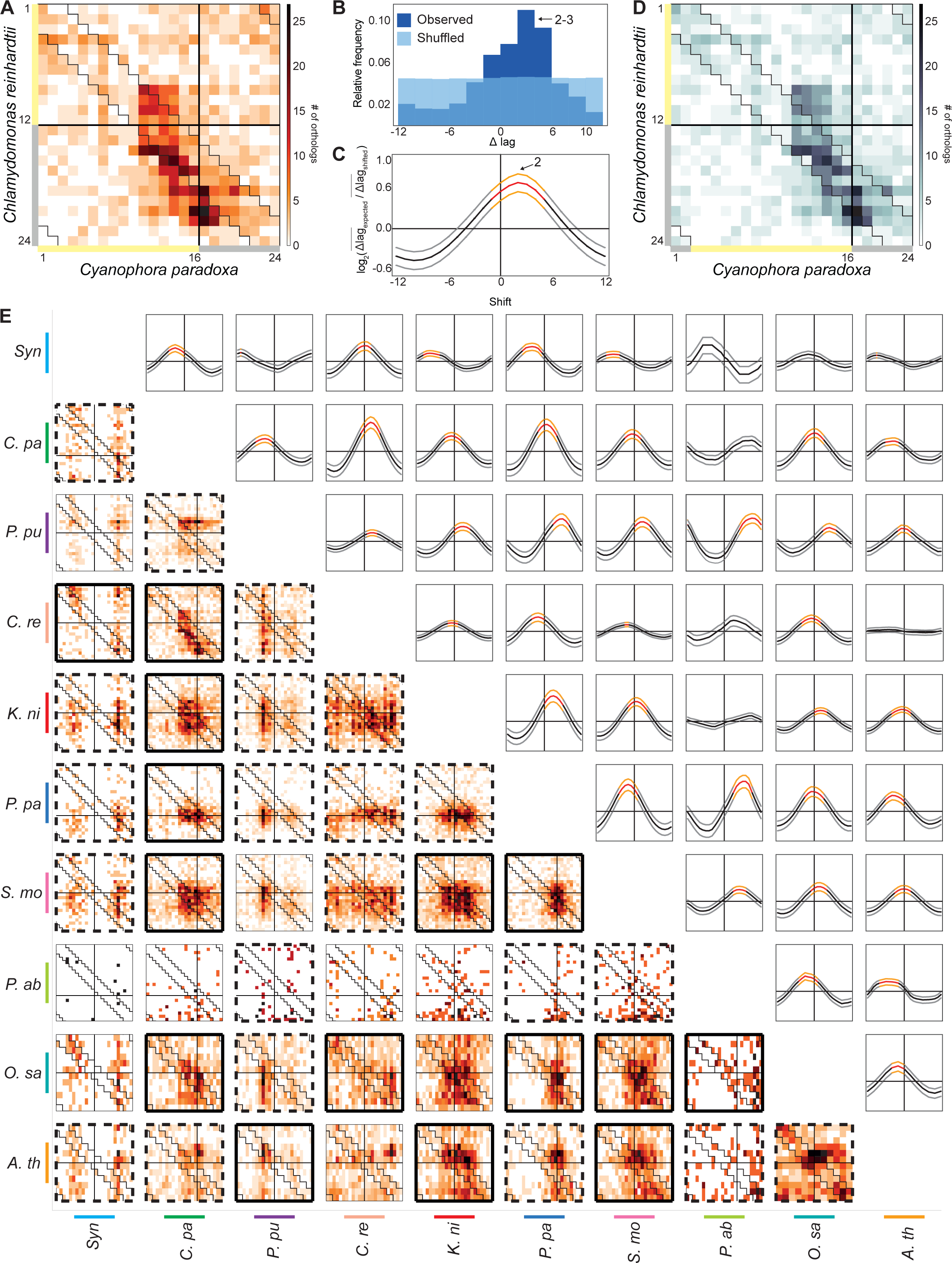
Comparison of diurnal transcriptomes. A) Comparison of lags of orthologs of *C. reinhardtii* (y-axis) and *C. paradoxa* (x-axis). The colour intensity of the cells indicates the number of orthologs that peak at a given lag combination in the two species. Black lines indicate a transition from light to dark, while the zigzag lines indicate lag differences within ±Δ2hours between the two species. B) Relative frequency of lag differences between orthologs (Δlag_observed_, dark blue bars, Δlag_expected_, light blue bars) in *C. reinhardtii* and *C. paradoxa*. Δlag values are binned in twos, where the first bar indicates [-12,-11], the second bar [-10,-9] and so on. C) Influence of the shift on transcriptome similarity. The middle thick line indicate the median of the log_2_(Δlag_expected_/Δlag_shifted_) values, whereas the upper and lower lines indicate the third and first quartiles, for every shift of *C. paradoxa* lag values within the interval of [-12, 12]. D) Comparison of lags of orthologs after the shift of +2 hours is applied to *C. paradoxa*. E) Comparison of all possible species combinations (heatmaps). The thick black frames of the heatmaps indicate which species combination has a significantly different Δlag_observed_ (FDR corrected empirical p-value < 0.05) from the Δlag_expected_, the dashed thick frames indicate which Δlag_observed_ is significantly different (FDR corrected empirical p-value < 0.05) than the Δlag_expected_ after the shift. The thin frames indicate combinations whose distribution is not significantly different than the permuted distribution.

### Diurnal transcriptomes are similar

To determine if the diurnal transcriptomes of the studied species are similar, we investigated whether orthologous genes peak at a similar time during the day in the ten species. To this end, we first identified orthologs among the species through protein sequence similarity analysis using OrthoFinder (Emms and Kelly, 2015). To avoid analysing unclear orthologous relationships, which can arise by gene duplications, we only considered orthologs which showed a one-to-one relationship.

We visualized similarities across two transcriptomes with a lag-lag heat map, which indicates when the orthologs in two species peaks during the day (Fig. 3A, an example from the Chlamydomonas-Cyanophora comparison). The lag-lag heat map would show two similar transcriptomes as points found in a diagonal that starts in the upper-left corner and ends in the lower-right corner (see an example of similar transcriptomes in Supplementary Fig. 4). The comparison between Chlamydomonas (12L/12D) and Cyanophora (16L/8D) shows that, despite the four hour difference in the photoperiod in which the algae were grown, most of the orthologs of these two species tend to be found on the diagonal of the lag-lag plot, indicating that the orthologs of these two species peaks at a similar time of the day.

**Fig. 4.**
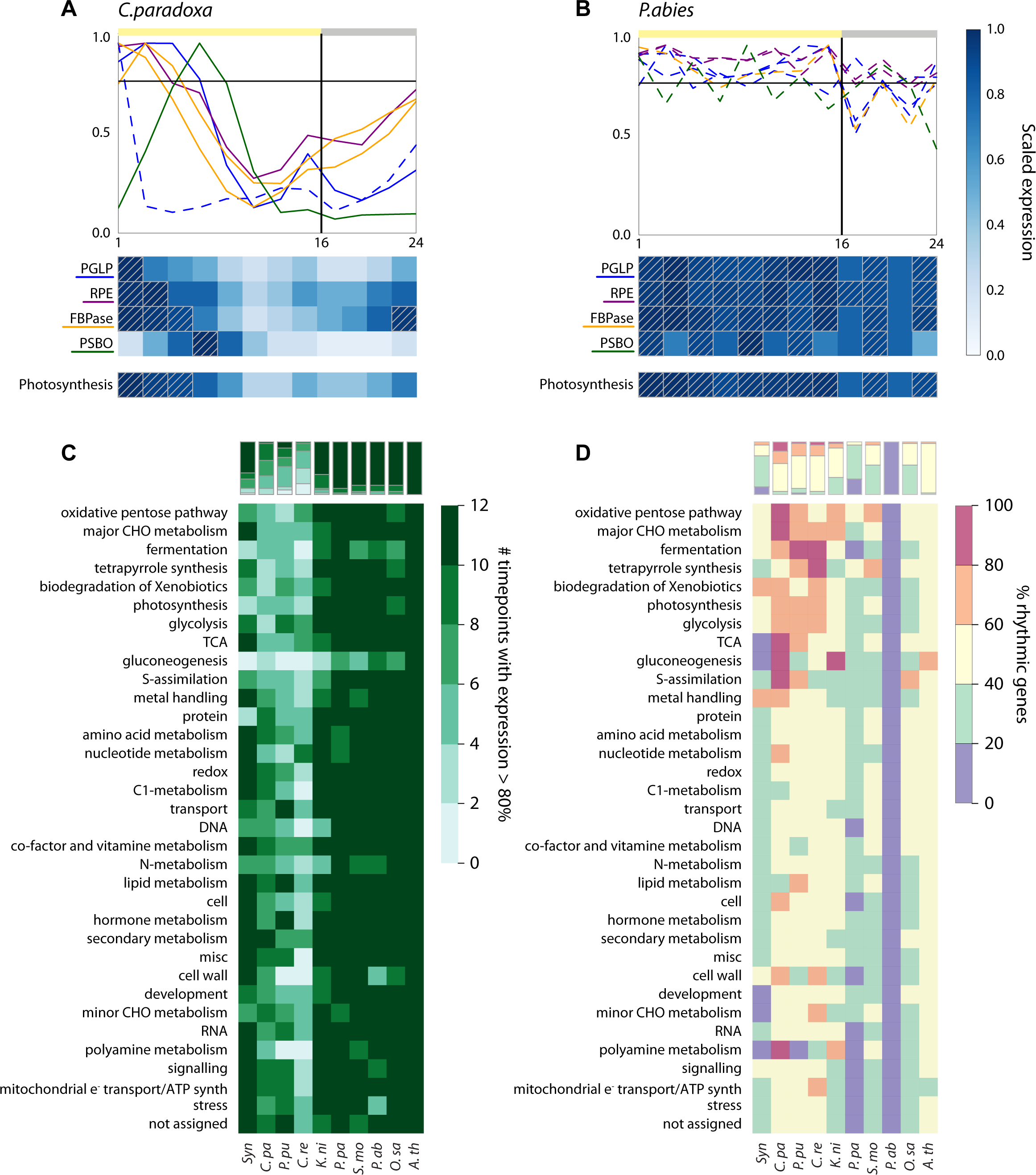
Diurnal expression of biological processes. A) Gene expression profiles of Cyanophora genes belonging to four families involved in photosynthesis. Solid lines indicate significantly rhythmic, while dashed lines indicate arrhythmic genes. The horizontal black line indicates the 80% expression threshold. The heat map below the gene expression plot shows the average expression of the four gene families and of all genes assigned to photosynthesis. Dashed squares indicate the average expression values higher than 80% at a given time point. B) Gene expression profiles of photosynthetic genes in spruce. C) Expression specificity of biological processes. Rows indicate MapMan bins while columns indicate the species. The colour intensity of the cells reflects the specificity of process expression, i.e. the number of timepoints where a bin is expressed above the average 80% of the maximum expression of the genes. D) Percentages of rhythmic genes per biological process per species. The amounts are adjusted to ranges (<=20%, between 20 and 40%, between 40 and 60%, between 60 and 80% and >80%). The bars above both heatmaps summarize the processes expression.

To estimate if diurnal transcriptome time-series are significantly similar in two species, we calculated the lag differences (Δlag) (i.e. differences in peaking times) between the orthologs (Fig. 3B, dark blue bars), expecting that the average lag differences of significantly similar transcriptomes would be smaller than expected by chance. To this end, we compared the average observed lag differences (Δlag_observed_) with the average lag differences calculated from permuted expression data (Δlag_expected_). Indeed, the analysis revealed that, in contrast to evenly distributed Δlag_expected_ values and Δlag_expected_ = 5.98, the Δlag_observed_ values peaked at +2 hours and Δlag_observed_ = 4.07 (Fig. 3B). Furthermore, comparison of the Δlag_observed_ and the Δlag_expected_ values revealed that the diurnal transcriptomes of Chlamydomonas and Cyanophora are significantly similar (FDR corrected empirical p-value < 0.012, obtained from 1000 permutations of expression data).

Interestingly, because Δlag_observed_ values peak at +2 hours when comparing Cyanophora and Chlamydomonas, this indicates that Cyanophora genes tend to peak two hours earlier than the corresponding Chlamydomonas genes. Consequently, the two transcriptomes would be more similar (i.e. Δlag_observed_ closer to 0) if the lags of Cyanophora genes were shifted forward by two hours. To test this, we calculated Δlag_expected_/Δlag_observed_ value between Cyanophora and Chlamydomonas, upon shifting the lag of Cyanophora genes by an integer in the interval of [-12, 12]. The value becomes larger than one in cases where the observed lag difference (Δlag_observed_) is smaller (i.e., more similar) than the lag differences obtained from the permuted expression data (Δlag_expected_). Indeed, we observed highest Δlag_expected_/Δlag_observed_ value when a shift of +2 is applied to Cyanophora transcriptome (Fig. 3C). Accordingly, the diagonal pattern in the lag-lag plot became more evident after the shift (Fig. 3D), which was accompanied by a more significant *p*-value (Supplementary Fig. 5). Thus, we conclude that the diurnal transcriptomes of Cyanophora and Chlamydomonas are significantly similar, but there is on average, a two hour difference between when the orthologs in these two organisms peak.

**Fig. 5.**
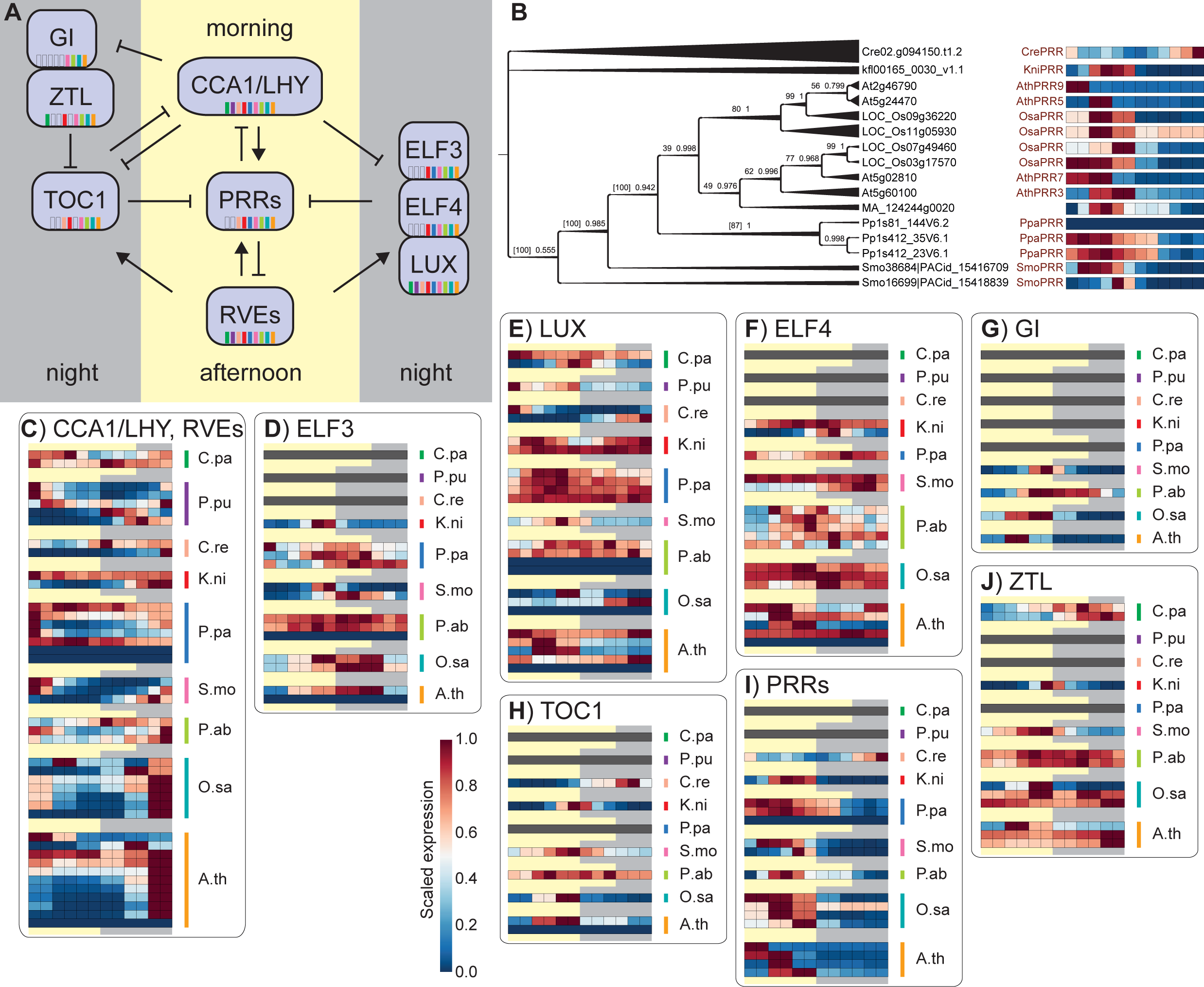
Analysis of clock core component genes. A) Simplified Arabidopsis circadian clock model (modified from (Hsu and Harmer, 2014)). B) Phylogenetic tree constructed on Bayesian inference of phylogeny. The first and the second number at the nodes indicate bootstrap values, and posterior probabilities, respectively. The gene annotation was based on (Linde *et al.*, 2017). The single heatmaps represent the 0-1-scaled expression profiles of each gene of the *PRR* family. Scaled expression profiles of genes belonging to C) *CCA1*/*LHY*/*RVE*, D) *ELF3*, E) *LUX*, F) *ELF4*, G) *GI*, H) *TOC1*, I) *PRRs*, and J) *ZTL*.

To investigate similarities of the transcriptomes across the Archaeplastida, we applied this analysis for all possible species combinations (Fig. 3E), and we observed that several comparisons show significant similarity even without applying a shift (solid thick frame, FDR corrected empirical *p*-value < 0.05) and most comparisons reach a significant similarity when the shift to their lags is applied (dashed thick frame, FDR corrected empirical *p*-value < 0.05; see Supplementary Fig. 5 upper left triangle for the shift applied in each case). Spruce continues to be the exception, being similar to only five species. Furthermore, spruce produces the most extreme shift values required to obtain significantly similar transcriptomes, as exemplified by comparison with Selaginella (shift of 5 hours), Physcomitrella (6 hours) and Porphyridium (9 hours, Supplementary Fig. 5).

We also investigated whether a specific event, such as colonization of land, led to reprogramming of diurnal gene expression. Such reprogramming would result in land plants having more similar diurnal transcriptome time-series than aquatic algae, which should result in smaller Δlag_expected_ /Δlag_observed,_ shift and p-values when comparing land plants. However, we could not detect any tendencies or specific patterns suggesting such reprogramming, as e.g., the Arabidopsis transcriptome was more similar to the aquatic Klebsormidium, than rice (Supplementary Fig. 6). This suggests that whereas the diurnal transcriptomes are similar, no dramatic reprogramming occurred at a specific time in plant evolution. These results also underline the point that there is variation even between more closely related species (e.g., Arabidopsis and rice), making it problematic to draw conclusions about long term changes from comparison of single-celled algae and plants.

### Evolution of diurnal expression of biological processes

To further compare the diurnal gene expression of the Archaeplastida, we investigated the expression of the major cellular pathways of every species. To this end, we analysed the expression profiles and the average expression of genes assigned to MapMan ontology bins (Thimm *et al.*, 2004). The analysis revealed that specific organisms show stronger rhythmicity for specific processes.

For example, most of the photosynthesis-related genes are rhythmic in Cyanophora, with highest expression during the light period, with an anticipatory decrease below 50% of maximum expression before dusk and anticipatory increase before the approaching dawn (Fig. 4A). Conversely, the corresponding orthologs in spruce are not rhythmic and expressed within 80% of maximum levels at most time points of the day (Fig. 4B, dashed squares). The average expression of the photosynthesis genes is summarized by the ‘photosynthesis’ heatmap which showed that photosynthesis genes are on average above 80% of their maximum expression levels in three and ten out of twelve time points for Cyanophora and spruce, respectively (Fig. 4A-B, dashed squares). This result indicates that spruce, in contrast to Cyanophora, does not rely on diurnal control of expression of photosynthesis genes.

To search for similarities and differences in diurnal expression of biological processes in Archaeplastida at a more general level, we performed this analysis for all MapMan bins (Fig. 4C). In line with the observation that cell division is not controlled by light in land plants (Fig. 1D), genes belonging to ‘cell’ bin (which contains ‘cell.division’ child bin) in multicellular organisms have a more ubiquitous expression compared to the defined expression of unicellular plants (Fig. 4C, ubiquitously expressed genes indicated by dark green colour). Furthermore, we observed that biological processes in single-celled algae, such as Cyanophora, Porphyridium, and Chlamydomonas tend to be expressed at specific times during the day (Fig. 4C, genes belonging to specifically expressed processes are indicated by light green cells). Conversely, in the relatively simple multicellular Klebsormidium and in land plants, biological processes show a more ubiquitous pattern of diurnal gene expression (Fig. 4C, dark green cells). This suggests that establishment of multicellularity, rather than colonization of land, decreased the diurnal regulation of gene expression. Nevertheless, up to 89% of the Arabidopsis transcriptome was shown to show rhythmicity when temperature was combined with the light/dark cycles (Michael *et al.*, 2008), suggesting that Arabidopsis (and likely all land plants) show an increased sensitivity to other external stimuli, not investigated in this study.

Finally, we examined how the rhythmicity of biological processes differs between species (Fig. 4D). We observed that in single-celled organisms, basal metabolic processes, such as the oxidative pentose pathway, major carbohydrate metabolism, fermentation, tetrapyrrole synthesis, biodegradation of xenobiotics, photosynthesis, glycolysis and tricarboxylic acid (TCA) cycle are the most rhythmic, with between 60 to 100% of the genes assigned to the processes showing significant oscillatory behaviour. Conversely, we observed a decrease of rhythmicity of most biological processes in Klebsormidium and land plants, with spruce showing less than 20% of rhythmic genes for all the investigated processes.

### Analysis of circadian clock components

The circadian clock has been extensively studied in Arabidopsis (Harmer *et al.*, 2000; Huang *et al.*, 2012; Hsu and Harmer, 2014) and it is known to regulate diurnal expression of thousands of transcripts (Harmer *et al.*, 2000; Covington *et al.*, 2008) to coordinate diverse biological processes so that they occur at the most appropriate season or time of day. The clock consists of 3 major, integrated feedback loops (Fig. 5A). A morning loop where *CIRCADIAN CLOCK ASSOCIATED 1* (*CCA1*) and *LATE ELONGATED HYPOCOTYL* (*LHY*) together positively regulate the expression of *PSEUDO RESPONSE REGULATOR* (*PRR*) genes, whereas the latter in response, negatively regulates the two genes (Gendron *et al.*, 2012; Nakamichi *et al.*, 2010; Kamioka *et al.*, 2016). The central loop has as protagonists *CCA1*/*LHY* and *TIME OF CAB EXPRESSION 1* (*TOC1*, also known as *PRR1*) which negatively regulate one another (Alabadí *et al.*, 2001). The last loop includes the evening complex, activated by the *REVEILLE* genes, formed by *LUX ARRHYTHMO* (*LUX*), *EARLY FLOWERING 3* and *4* (*ELF3*, *ELF4*), that together with *GIGANTEA* (*GI*) and *ZEITLUPE* (*ZTL*) proteins, represses *TOC1* and the other members of *PRR* ensuring the expression of the morning-phased components *CCA1* and *LHY* (Rawat *et al.*, 2011; Farinas and Mas, 2011; Nusinow *et al.*, 2011; Miyazaki *et al.*, 2015).

To explore how phototropism and photoperiodism evolved we analysed the expression of eight gene families containing the core clock genes from Arabidopsis (*CCA1*, *LHY*, *RVE1-8*, *PRR3*, *5*, *7*, *9*, *TOC1*, *LUX*, *ELF3*, *ELF4*, *GI* and *ZTL*) in the nine members of the Archaeplastida (Fig. 5C-J). As previously shown (Serrano-Bueno *et al.*, 2017), only *CCA1*/*LHY*/*RVE* and *LUX* families are present in most of the species, indicating that the three major loops are not complete in the early-diverging algae Cyanophora and Porphyridium, because *PRR* genes (morning loop), *TOC1* (central loop), *ELF3*/*ELF4*/*GI*/*ZTL* (evening loop) are absent in these two algae (Fig. 5A, C-J). These results suggest that despite the incomplete Viridiplantae clock, conserved diurnal gene expression can be mediated by other mechanisms, such as light sensor-mediated response to light (de los Reyes *et al.*, 2017; Serrano-Bueno *et al.*, 2017), carbon signalling (Blasing, 2005; B. Usadel *et al.*, 2008; Flis *et al.*, 2016), or an uncharacterized type of clock. Furthermore, the different loops were established at specific times in plant evolution, because genes constituting the morning and central loop are first found in the Viridiplantae clade (containing Chlamydomonas, Klebsormidium and land plants, Supplementary Fig. 8-9), whereas a complete evening loop was established in streptophytes (containing Klebsormidium and land plants, Supplementary Fig. 11-12).

Next, we compared expression patterns of the clock genes (Fig. 5C-J). The *PRR* genes show conserved expression during the day across all species (Fig. 5I, Supplementary Fig. 8), with the exception of Chlamydomonas *PRR*, which is mostly expressed during the night. A similar behaviour was observed for *TOC1*, whereby all species showed the highest expression of the gene at dusk, whereas Chlamydomonas *TOC1* peaked at night, confirming previous analyses (Mittag *et al.*, 2005; de los Reyes *et al.*, 2017). The larger gene families exhibited more complex expression patterns. *CCA1*/*LHY*/*RVEs* was expressed predominantly at the end of night/beginning of the day in all the species (Fig. 5C), whereas *ELF3* showed a different expression at the end of the day/beginning of the night (Fig. 5D).

Our analyses reveal that spruce has the smallest fraction of rhythmic genes, with only 5.2% genes showing significantly rhythmic behaviour (Fig. 1B). Surprisingly, all of the clock components are present in spruce (Fig. 5C-J), suggesting that the clock is running differently or is not active in spruce. We observed that *ELF3* (Fig. 5D), *GI* (Fig. 5G) and *TOC1* (Fig. 5H), which are typically expressed at a specific time point in the other species, are expressed more broadly in spruce. Whereas RNA sequencing data, due to providing only relative gene expression abundances, cannot reveal whether the clock is not active, the lower specific expression pattern of these key genes might explain the low frequency of rhythmic transcripts in the spruce diurnal transcriptome.

## Conclusion

The life of photosensing organisms is finely tuned by photoperiodism, temperature, availability of photosynthetic products, and the circadian clock (Lagercrantz, 2009; B. Usadel *et al.*, 2008; Björn Usadel *et al.*, 2008; Nohales and Kay, 2016). Our findings show that most members of the Archaeplastida subject more than one-third of their genes to diurnal gene expression (Fig. 1B), underlining the importance of diurnal regulation. We demonstrated that, whereas the age of a gene family positively correlates with the level of rhythmicity and expression of genes (Fig. 2C-D), it does not affect diurnal expression patterns (Fig. 2A, Supplementary Fig. 2). This suggests that gene expression levels and expression patterns are controlled by two separate mechanisms.

We found that diurnal programs are remarkably similar, despite the large phylogenetic distance spanning more than 1.5 billion years of evolution, loss of light-controlled cell division in land plants (Fig. 1D), and wide morphological diversity of the analysed species (Fig. 3E). Interestingly, diurnally-regulated orthologs tend to be expressed at approximately the same time in the 24 h light-dark cycle, although the precise expression peak can shift by a few hours (Fig. 3E). This leads us to conclude that the sequence of the diurnal transcriptomes is similar because such an arrangement presents an optimal sequence of gene expression, regardless of morphology and habitat and was established early in Archaeplastida evolution. The exception to this rule is spruce which is arrhythmic, likely due to adaptation to dramatic changes in day length (21 hours daylight in July and 4 hours in December in Umeå, Sweden). This suggests that despite its universality and ancient origin, diurnal gene expression can be rewired to better fit the local environment.

We observed that unicellular organisms exhibit a more specific diurnal gene expression, often restricting the expression of the major biological pathways to an eight hour period (Fig. 4C), and with a higher rhythmicity of basal processes, such as photosynthesis, major carbohydrate metabolism, oxidative pentose pathway, and tetrapyrrole synthesis. Conversely, Klebsormidium and land plants exhibit a more uniform expression of biological processes. We hypothesize that the increased complexity of multicellular organisms necessitated an uncoupling of the diurnal regulation of biological processes, specifically cell division, from the external influence of the light/dark alternation. Furthermore, we cannot exclude that the combination of signals from the diurnal transcriptomes of the different cell types obscures the specific diurnal expression profiles of, e.g., epidermis and mesophyll cells (Thain *et al.*, 2000; Yakir *et al.*, 2011; Gould *et al.*, 2018). However, the Klebsormidium data provide strong evidence for light-controlled cell division (Fig. 1D), in this species that possesses only a single cell type, but similar to land plants, shows a more uniform expression of biological processes. This result suggests that the more ubiquitous expression pattern evolved in streptophytes.

Our analysis confirms the ancient origin of the circadian clock by identifying *CCA1*-and *LUX*-like genes in glaucophytes and rhodophytes (Fig. 5B, Supplementary Fig. 7-14). However, the other essential components of the three main control loops are absent in these two species, suggesting that their clock might function differently from the well-studied clock in streptophytes. Additional studies, with experimental setups that minimize the influence of cell division and photoperiodism are needed to understand how the clock functions in these anciently diverged algae.

The observed conservation of the diurnal transcriptomes underlines how critical light sensing is in all life forms. Investigating if the conservation of the diurnal regulation is also reflected by conserved oscillation of proteins and metabolites will be an interesting area of future investigation.

## Methods

### Growth conditions

*Cyanophora paradoxa* UTEX555 (SAG 29.80, CCMP329) was obtained from Provasoli-Guillard National Center for Marine Algae and Microbiota (NCMA, Bigelow Laboratory for Ocean Sciences). The cultures were grown in C medium (Ichimura, 1971), at 24°C in 16L/8D photoperiod (40 μmol photons m^-2^ s^-1^), aerated with normal air. *Porphyridium purpureum* SAG 1380-1d was obtained from SAG (Culture Collection of Algae at Göttingen University). The cultures were grown in ASW medium (KESTER *et al.*, 1967), at 25°C in 12L/12D photoperiod (100 μmol photons m^-2^ s^-1^). *Klebsormidium nitens* was obtained from the NIES collection (Tsukuba, Japan). The cultures were grown in liquid C medium at 24°C, in 12L/12D photoperiod (40 μmol photons m^-2^ s^-1^). *Physcomitrella patens* was grown in Jiffy −7 peat pellets at 23°C, in 16L/8D photoperiod (100 μmol photons m^-2^ s^-1^) and the gametophytes were collected. *Selaginella moellendorffii* was grown on modified Hoagland solution (0.063 mM FeEDTA; 0.5 mM KH_2_PO_4_; 2.5 mM KNO_3_; 2 mM CA(NO_3_)_2_x4H_2_O; 1 mM MgSO_4_x7H_2_O; 50.14 µM H_3_BO_3_; 9.25 µM MnCl_2_x4H_2_O; 1 µM ZnCl_2_; 1 µM CuCl_2_; 0.5 µM Na_2_MoO_4_x2H_2_O) at 24°C, in 12L/12D photoperiod (70 μmol photons m^-2^ s^-1^) and microphylls were collected. All organisms were actively growing. *Picea abies* seedlings were grown in soil at 24°C, in 16L/8D photoperiod (250 μmol photons m^-2^ s^-1^) and needles were collected.

### RNA isolation and RNA sequencing

The total RNA was extracted using Spectrum™ Plant Total RNA Kit (Sigma-Aldrich) according to the manufacturer’s instructions. The integrity and concentration of RNA was measured using RNA nano chip on Agilent Bioanalyzer 2100. The libraries were prepared from total RNA using polyA enrichment and sequenced using Illumina-HiSeq2500/4000 at Beijing Genomics Institute and Max Planck-Genome-centre in Cologne.

## Data availability

The TPM-normalized expression data is available in Supplemental Table 2-11, while the fastq files representing the raw RNA sequencing data are available from EBI Array Express under accession number E-MTAB-.

### Analysis of RNAseq data and microarrays data

The reads were trimmed, mapped, counted and TPM-normalized using the LSTrAP pipeline (Proost *et al.*, 2017). The genomes used for mapping are for Cyanophora v1.0 (the unpublished genome update obtained from Dana Price, SUNJ), Porphyridium v1.0 (obtained from *The Porphyridium purpureum Genome Project*)(Bhattacharya *et al.*, 2013), Klebsormidium v1.0 (obtained from *Klebsormidium nitens* NIES-2285 genome project)(Hori *et al.*, 2014), Picea v1.0 (obtained from PlantGenIE)(Nystedt *et al.*, 2013) and Physcomitrella v1.6 (obtained from Cosmoss)(Rensing *et al.*, 2008). Reads from Selaginella were mapped to the transcriptome. Chlamydomonas RNA-seq counts of samples taken every two hours, obtained from (Zones *et al.*, 2015) were TPM normalized. Synechocystis microarray raw data, obtained from ArrayExpress accession E-GEOD-47482 (Beck *et al.*, 2014) was processed using the R-package limma (Ritchie *et al.*, 2015), while Arabidopsis raw data, obtained from E-GEOD-3416 (Blasing, 2005) were RMA normalized. Processed rice expression data was obtained from E-GEOD-28124 (Xu *et al.*, 2011). The quality of the samples and agreement of the replicates was assessed through PCA analysis. For further analyses only expressed genes were selected from the normalized matrices. A gene was identified as expressed if the TPM value was > 1 for the RNAseq data or called ‘present’ at least twice (once) in at least one time point for Arabidopsis (Oryza).

### Detection of rhythmic genes

Rhythmic genes were identified by using the JTK algorithm (Hughes *et al.*, 2010). For Cyanophora, Porphyridium, Chlamydomonas, Klebsormidium, Physcomitrella, Selaginella, and Picea we used a total of 12 timepoints (ZT1 to ZT23), and the replicate number was set to 3 (2 for Chlamydomonas). The temporal spacing between samples was set to 2 hours and the target period was set to 12 h (effectively equal to 24h). The Synechocystis dataset consisted of one replicate per timepoint over two day cycle for a total of 24 timepoints, and the JTK parameters were set to: number of replicates to 1 and the target period 48 h (effectively equal to 24h). For Arabidopsis and Oryza the temporal spacing between samples was set to 4h and the target period to 6h (effectively equal to 24h). For Arabidopsis, the number of replicates was 3 while for Oryza 2. Using an adjusted *p*-value cutoff < 0.05 we identified 2440, 12343, 6597, 12341, 9375, 11692, 11860, 5280, 8664 and 6409 rhythmic genes for Synechocystis, Cyanophora, Porphyridium, Chlamydomonas, Klebsormidium, Physcomitrella, Selaginella, Picea, Oryza, and Arabidopsis, respectively (Supplementary Table 2-11). In order to calculate the percentages of rhythmic genes in a comparable manner we performed an additional run of JTK where the initial expression matrices were adapted to the dataset with the least number of timepoints and replicates (Oryza), therefore we set the number of replicates to 2, the spacing between samples to 4h and the target period was set to 6h (Supplementary Fig. 15). LAG 0 was identified as the first time point collected after the light was turned on (i.e., LAG 0 in Cyanophora was corrected to LAG 1, LAG 0 in Arabidopsis was corrected to LAG 4).

To validate the results obtained by JTK we used the Haystack algorithm (Mockler *et al.*, 2007)(Supplementary Fig. 16). The correlation cutoff was set to 0.7, the fold-change to 1, the background cutoff to 1 and a p-value cutoff of 0.05. The used five models to infer the rhythmicity. We detected 3004, 3756, 6354, 13429, 5630, 3059, 12914, 20613 and 12507 rhythmic genes for Synechocystis, Cyanophora, Porphyridium, Chlamydomonas, Klebsormidium, Physcomitrella, Selaginella, Picea, Oryza, and Arabidopsis, respectively (Supplementary Table 2-11). The analysis showed a good agreement between the significantly rhythmic genes, and their estimated lags (Supplementary Fig. 15).

### Detection of orthologous genes with Orthofinder

Orthologous genes were obtained by using OrthoFinder v1.1.8 (Emms and Kelly, 2015) with default parameters and Diamond (Buchfink *et al.*, 2015), which identified 12913 orthogroups (Supplementary Table 13). For further analysis, we considered only one-to-one orthologs, in order to ensure that the comparison between genes would dismiss comparisons between unclear orthologs.

### Phylostratigraphic analysis of rhythmicity and average expression

The phylostratum of each orthogroup was estimated by finding the oldest clade in the family. To estimate if a given phylostratum is significantly rhythmic, we first counted how many genes are rhythmic in a phylostratum. Next, we performed a permutation analysis where we shuffled the gene-phylostratum assignment 1000 times, and we obtained a number of rhythmic genes per phylostratum. The empirical p-value for enrichment was obtained by counting the number of times the permuted distribution showed a higher number of rhythmic genes compared to the observed values, divided by the number of permutations (1000). The empirical *p*-values for depletion were calculated in the same manner.

The gene average expression of the phylostrata was obtained from the average gene expression of each gene assigned to each phylostratum. To identify averages expression that are significantly different, we used a two-sample Kolmogorov–Smirnov test (K–S test) for all possible combinations. The obtained *p*-values were FDR corrected. The results of the comparison are found in Supplementary Fig. 3.

### Estimating the significance of similarity of diurnal transcriptomes

To estimate the similarity of diurnal transcriptomes of two species, we first calculated the absolute difference of expression (Δlag_observed_) among each pair of one-to-one orthologs. Because of the cyclic nature of the data, for *abs*(Δlag_observed_) > 12, we used *abs*(Δlag_observed_-24). Next, we calculated this number for all one-to-one orthologs to arrive at the average Δlag_observed_. Finally, to test whether a similar distribution of pairs would have been obtained by chance, we shuffled the columns (timepoints) in the expression matrices 1000 times, and ran JTK for each permutation. For each permutation, we calculated the Δlag value, as described above, to arrive at 1000 Δlag_expected_ values. The empirical *p*-value was calculated by counting how many times the Δlag_observed_ was smaller than the Δlag_expected_ values. The *p*-values were then FDR corrected.

The shift analysis was performed by adding an integer in the interval of [-12, 12] to the lag values all the rhythmic genes of the older species, and calculating Δlag_observed_ upon the shift (here defined as Δlag_shifted_). For each shift, the empirical *p*-value was calculated, as described above, and the log_2_(Δlag _expected_/Δlag_shifted_) was calculated. The first quartile, the median and the third quartile of the distribution was plotted and color coded according to indicate significantly (orange, red) and not significantly (grey, black) similar transcriptome upon the shift. The shift which provided the highest log_2_(Δlag_expected_/Δlag_shifted_) was used to identify the lowest p-value (Figure 3C). The *p*-values corresponding to the best shift for each species comparison were then FDR corrected (Figure 3E).

### Diurnal expression of biological processes

We used Mercator with standard settings to annotate the coding sequence with MapMan bins (Lohse *et al.*, 2014). To investigate the expression of the biological processes, we analyzed the more general, first level MapMan bins. For each gene in a bin, expression vector of the gene was first scaled by dividing the vector by the maximum value of the vector. Then, the expression vectors of the genes within the bin were summed, and the summed vector was divided by the maximum value. Finally, to calculate the specificity of the MapMan bin expression, we counted the number of time points with value > 0.8.

### Rhythmicity of biological processes

The rhythmicity of the processes was studied by calculating the percentage of rhythmic genes assigned to a given first level MapMan bin. To make the results comparable between the 4 hours (Arabidopsis and rice) and 2 hours experiments (every other Plantae member), we removed every other time point from the 2 hours expression data, to arrive at 4 hours expression matrices. The expression matrices were then analysed by JTK to arrive at rhythmic genes, which were used to generate Figure 4D.

### Phylogenetic analysis and tree construction

To identify functionally related genes of the circadian clock in the nine species, we selected eight orthogroups (OG0000138, OG0000233, OG0001151, OG0001171, OG0001580, OG0003526, OG0007425, OG0008362) containing the 20 major clock components from *Arabidopsis thaliana*. The protein sequences were aligned using MUSCLE software (Edgar, 2004) and trimmed with TrimAl with standard settings (Capella-Gutiérrez *et al.*, 2009). The best-fit models for each orthogroups were identified using ProtTest 3.4.2 (Darriba *et al.*, 2011), which was based on the Bayesian information criterion (BIC). The phylogenetic reconstruction was done by analyzing the protein alignments with both Maximum Likelihood (ML) and Bayesian inference (BI) methods. The ML analysis was performed using PhyML 3.0 algorithm (Guindon *et al.*, 2010)with a bootstrap with 100 iterations. Bayesian analysis was conducted using MrBayes 3.2.6 (Ronquist *et al.*, 2012) where two Markov Chain Monte Carlo (MCMC) runs were performed for 1 million generations, and sampling was performed every 500 generations. Both the average standard deviation of split frequencies and potential scale reduction factor (PSRF) were < 0.01 and close to 1, respectively, across the two runs. The trees were visualized using TreeGraph2 (Stöver and Müller, 2010) using the Bayesian tree as a reference. The bootstrap values from PhyML and the posterior probabilities values from MrBayes were mapped as branch support. For large orthogroups, such as PRRs (OG0000138), *CCA1*/*LHY-RVEs* (OG0000233) and ZTL (OG0001151), we isolated the relevant clades by analysing phylogenetic trees based on Maximum Likelihood (ML) and Bayesian inference (BI) methods (Supplementary Fig. 7-14), and the phylogenetic analysis was run on the selected genes. As previously reported (Linde *et al.*, 2017), we could not separate *CCA1*/*LHY* from *RVE* genes since both classes of proteins are MYB-like transcription factors (Wang, 1997; Schaffer *et al.*, 1998; Rawat *et al.*, 2009)(Supplementary Fig. 7).

---

**Supplementary Fig. 1. PCA analysis of the expression data.** Each dot represents a sample. The dots are color-coded based on the time of the sampling, with dark red being the first sample after the light was turned on.

**Supplementary Fig. 2. Distribution of expression peaks of rhythmic genes.** The genes are divided by the phylostrata (rows), and the expression peaks are plotted through the lags (columns). The color of the cells indicates the 0-1-scaled gene expression values, where 0 and 1 indicate the lowest and highest expression, respectively.

**Supplementary Fig. 3. Differential analysis of average expression of phylostrata.** The cell colors indicate the two-sample Kolmogorov–Smirnov comparison p-value (K–S test, FDR corrected) for all possible phylostrata combinations for average expression distribution.

**Supplementary Fig. 4. Comparison diurnal transcriptomes of two independent Arabidopsis experiments.** The color intensity of the cells indicates the number of genes that peak at a given lag combination in the two experiments. Black lines indicate a transition from light to dark, while the zigzag lines indicate an interval of 4 hours.

**Supplementary Fig. 5. Comparison of shifted diurnal transcriptomes.** The heatmaps indicate the comparisons of lags of orthologs of all possible species combination after the shift is applied to one of the two species (x-axis). The color intensity of the cells indicates the number of orthologs that peak at a given lag combination in the two species. Black lines indicate a transition from light to dark, while the zigzag lines indicate lag differences within ±Δ2hours between the two species. The thick black frames of the heatmaps indicate which species combination has a significantly similar Δlag_expected_ value (FDR corrected empirical p-value < 0.05). The thin frames indicate combinations which are not significantly similar. The upper right triangle show, from top to bottom, the number of orthologs included in the analysis, the shift and the p-value (FDR corrected), respectively.

**Supplementary Fig. 6. Analysis of reprogramming events.** A) FDR-corrected empirical *p*-value distribution signifying the similarities of the transcriptomes. B) Shift values producing the highest Δlag_expected_/Δlag_observed_ values. C) Distribution of the highest Δlag_expected_/Δlag_observed_ obtained by the best shift.

**Supplementary Fig. 7. Inferred phylogenetic tree for CCA1/LHY/RVEs.** The phylogenetic tree was constructed based on bayesian inference of phylogeny, and the bootstrap values were estimated on Maximum-Likelihood phylogeny of the orthogroup containing Arabidopsis CCA1/LHY and RVE 1-8. The gene annotation was based on (Linde *et al.*, 2017). The single heatmaps represent the 0-1-scaled expression profiles of each gene.

**Supplementary Fig. 8. Inferred phylogenetic tree for PRRs.** The phylogenetic trees were constructed based on the identification of the orthogroup containing Arabidopsis PRR3, 7, 5, 9.

**Supplementary Fig. 9. Inferred phylogenetic tree for TOC1.** The phylogenetic trees were constructed based on the identification of the orthogroup containing Arabidopsis TOC1.

**Supplementary Fig. 10. Inferred phylogenetic tree for LUX.** The phylogenetic trees were constructed based on the identification of the orthogroup containing Arabidopsis LUX.

**Supplementary Fig. 11. Inferred phylogenetic tree for ELF3.** The phylogenetic trees were constructed based on the identification of the orthogroup containing Arabidopsis ELF3.

**Supplementary Fig. 12. Inferred phylogenetic tree for ELF4.** The phylogenetic trees were constructed based on the identification of the orthogroup containing Arabidopsis ELF4 gene.

**Supplementary Fig. 13 Inferred phylogenetic tree for ZTL.** The phylogenetic trees were constructed based on the identification of the orthogroup containing Arabidopsis ZTL.

**Supplementary Fig. 14. Inferred phylogenetic tree for GI.** The phylogenetic trees were constructed based on the identification of the orthogroup containing Arabidopsis GI.

**Supplementary Fig. 15. Influence of sample density on the percentage of detected rhythmic genes.** The x-and y-axis show the species and the percentage of rhythmic genes in a species, at a given dataset. Bars represent percentages of rhythmic genes identified by JTK algorithm. Dark blue bars are calculated on the original datasets (sampled every 2 hours, in 3 replicates, 2 for Synechocystis and Chlamydomonas), light blue bars represent same datasets, but with 2 replicates, 6 time points and starting at ZT1, while white bars represent the datasets with 2 replicates, 6 time points starting at ZT3, the gray bar is calculated on the Arabidopsis adapted dataset with 2 replicates.

**Supplementary Fig. 16. Comparison of JTK and Haystack algorithms for rhythmic genes detection.** A) Venn Diagrams representing genes identified as rhythmic by JTK (green) and Haystack (blue). B) Comparison of lags of rhythmic genes identified by JTK (y-axis) and Haystack (x-axis). C) Stacked bars showing the percentages of rhythmic genes assigned to the same lag (dark red), to lags whose Δlag <=2 (pink) or Δlag>2 (light blue) by the two algorithms.

**Supplemental Table 1. LSTrAP results.** Percentages of reads mapped to the genome, counts of reads mapped to coding genes (“mapped”), mapped to non coding genes (“no-feature”) and not uniquely (“ambiguous”) with relative percentages.

**Supplemental Table 2-11. Species Matrices.** Each table shows: gene name, phylostratum, JTK adjusted p-value, JTK lag, mercator annotation and TPM (RMA) diurnal expression values for Synechocystis (2), Cyanophora (3), Porphyridium (4), Chlamydomonas (5), Klebsormidium (6), Physcomitrella (7), Selaginella (8), spruce (9), rice (10) and Arabidopsis (11).

**Supplemental Table 12. Age of gene families.** The first column indicates the oldest and the youngest species present in the families, while the second column contains the respective gene families.

**Supplemental Table 13. OrthoFinder output containing the gene families.**

## References

Alabadí, D., Oyama, T., Yanovsky, M.J., Harmon, F.G., Más, P. and Kay, S.A. (2001) Reciprocal regulation between TOC1 and LHY/CCA1 within the Arabidopsis circadian clock. Science (80-. )., 293, 880–883.

Archer, S.N., Laing, E.E., Möller-Levet, C.S., et al. (2014) Mistimed sleep disrupts circadian regulation of the human transcriptome. Proc. Natl. Acad. Sci., 111, E682–E691. Available at: http://www.pnas.org/lookup/doi/10.1073/pnas.1316335111.

Beck, C., Hertel, S., Rediger, A., et al. (2014) Daily expression pattern of protein-encoding genes and small noncoding RNAs in synechocystis sp. strain PCC 6803. Appl. Environ. Microbiol. ,80, 5195–5206.

Bell-Pedersen, D., Cassone, V.M., Earnest, D.J., Golden, S.S., Hardin, P.E., Thomas, T.L. and Zoran, M.J. (2005) Circadian rhythms from multiple oscillators: Lessons from diverse organisms. Nat. Rev. Genet., 6, 544–556.

Bhattacharya, D., Price, D.C., Chan, C.X., et al. (2013) Genome of the red alga Porphyridium purpureum. Nat. Commun., 4, 1941. Available at: http://www.pubmedcentral.nih.gov/articlerender.fcgi?artid=3709513&tool=pmcentrez&rendertype=abstract.

Blasing, O.E. (2005) Sugars and Circadian Regulation Make Major Contributions to the Global Regulation of Diurnal Gene Expression in Arabidopsis. Plant Cell Online, 17, 3257–3281. Available at: http://www.plantcell.org/cgi/doi/10.1105/tpc.105.035261.

Bowman, J.L., Floyd, S.K. and Sakakibara, K. (2007) Green genes-comparative genomics of the green branch of life. Cell, 129, 229–34. Available at: http://www.sciencedirect.com/science/article/pii/S0092867407004618.

Brinker, M., Wissel, K., Jäschke, K., Kellmann, J.W. and Piechulla, B. (2001) Circadian gene expression in angiosperms and gymnosperms. Int. J. Endoctyobiosis Cell Res., 14, 33–44.

Buchfink, B., Xie, C. and Huson, D.H. (2015) Fast and sensitive protein alignment using DIAMOND. Nat. Methods, 12, 59–60. Available at: http://www.nature.com/doifinder/10.1038/nmeth.3176%5Cnhttp://dx.doi.org/10.1038/nmeth.3176%5Cnhttp://www.nature.com/doifinder/10.1038/nmeth.3176%5Cn http://www.ncbi.nlm.nih.gov/pubmed/25402007.

Capella-Gutiérrez, S., Silla-Martínez, J.M. and Gabaldón, T. (2009) trimAl: A tool for automated alignment trimming in large-scale phylogenetic analyses. Bioinformatics, 25, 1972–1973.

Chang, C., Bowman, J.L. and Meyerowitz, E.M. (2016) Field Guide to Plant Model Systems. Cell, 167, 325–339.

Covington, M.F., Maloof, J.N., Straume, M., Kay, S.A. and Harmer, S.L. (2008) Global transcriptome analysis reveals circadian regulation of key pathways in plant growth and development. Genome Biol., 9.

Darriba, D., Taboada, G.L., Doallo, R. and Posada, D. (2011) ProtTest-HPC: Fast selection of best-fit models of protein evolution. In Lecture Notes in Computer Science (including subseries Lecture Notes in Artificial Intelligence and Lecture Notes in Bioinformatics). pp. 177–184.

Dodd, A.N. (2005) Plant Circadian Clocks Increase Photosynthesis, Growth, Survival, and Competitive Advantage. Science (80-. )., 309, 630–633. Available at: http://www.sciencemag.org/cgi/doi/10.1126/science.1115581.

Dormling, I., Gustafsson, Å. and Wettstein, D. von (1968) The experimental control of the life cycle in Picea abies (L.) Karst. Some basic experiments on the vegetative cycle. Silvae Genet, 17, 44–64.

Edgar, R.C. (2004) MUSCLE: Multiple sequence alignment with high accuracy and high throughput. Nucleic Acids Res., 32, 1792–1797.

Emms, D.M. and Kelly, S. (2015) OrthoFinder: solving fundamental biases in whole genome comparisons dramatically improves orthogroup inference accuracy. Genome Biol., 16.

Farinas, B. and Mas, P. (2011) Histone acetylation and the circadian clock: A role for the MYB transcription factor RVE8/LCL5. Plant Signal. Behav., 6, 541–543.

Flis, A., Sulpice, R., Seaton, D.D., Ivakov, A.A., Liput, M., Abel, C., Millar, A.J. and Stitt, M. (2016) Photoperiod-dependent changes in the phase of core clock transcripts and global transcriptional outputs at dawn and dusk in Arabidopsis. Plant Cell Environ., 39, 1955–1981.

Gendron, J.M., Pruneda-Paz, J.L., Doherty, C.J., et al. (2012) Overlapping and Distinct Roles of PRR7 and PRR9 in the Arabidopsis Circadian Clock. Plant Cell, 23, 90–99. Available at: file://userfs/jar563/w2k/Downloads/Review Papers/PNAS-2010-Zhu-13960-5.pdf%5Cnfile:///C:/Users/jar563/AppData/Local/Mendeley Ltd./Mendeley Desktop/Downloaded/Herrero et al. - 2012 - EARLY FLOWERING4 recruitment of EARLY FLOWERING3 in the nucleus sustains the.

Gerstein, M.B., Rozowsky, J., Yan, K.-K., et al. (2014) Comparative analysis of the transcriptome across distant species. Nature, 512, 445–8.

Goodstein, D.M., Shu, S., Howson, R., et al. (2012) Phytozome: A comparative platform for green plant genomics. Nucleic Acids Res., 40.

Gould, P.D., Domijan, M., Greenwood, M., Tokuda, I.T., Rees, H., Kozma-Bognar, L., Hall, A.J. and Locke, J.C. (2018) Coordination of robust single cell rhythms in the Arabidopsis circadian clock via spatial waves of gene expression. Elife, 7, e31700. Available at: https://elifesciences.org/articles/31700.

Guindon, S., Dufayard, J.F., Lefort, V., Anisimova, M., Hordijk, W. and Gascuel, O. (2010) New algorithms and methods to estimate maximum-likelihood phylogenies: Assessing the performance of PhyML 3.0. Syst. Biol., 59, 307–321.

Guo, Y.-L. (2013) Gene family evolution in green plants with emphasis on the origination and evolution of Arabidopsis thaliana genes. Plant J., 73, 941–51.

Gyllenstrand, N., Karlgren, A., Clapham, D., Holm, K., Hall, A., Gould, P.D., Källman, T. and Lagercrantz, U. (2014) No time for spruce: Rapid dampening of circadian rhythms in picea abies (L. Karst). Plant Cell Physiol., 55, 535–550.

Harmer, S.L., Hogenesch, J.B., Straume, M., Chang, H.S., Han, B., Zhu, T., Wang, X., Kreps, J.A. and Kay, S.A. (2000) Orchestrated transcription of key pathways in Arabidopsis by the circadian clock. Science (80-. )., 290, 2110–2113.

Heide, O.M. (1974) Growth and Dormancy in Norway Spruce Ecotypes (Picea abies) I. Interaction of Photoperiod and Temperature. Physiol. Plant., 30, 1–12.

Hori, K., Maruyama, F., Fujisawa, T., et al. (2014) Klebsormidium flaccidum genome reveals primary factors for plant terrestrial adaptation. Nat. Commun., 5, 3978. Available at: http://www.nature.com/ncomms/2014/140528/ncomms4978/full/ncomms4978.html.

Hsu, P.Y. and Harmer, S.L. (2014) Wheels within wheels: The plant circadian system. Trends Plant Sci., 19, 240–249.

Huang, W., Pérez-García, P., Pokhilko, A., Millar, A.J., Antoshechkin, I., Riechmann, J.L. and Mas, P. (2012) Mapping the core of the Arabidopsis circadian clock defines the network structure of the oscillator. Science (80-. )., 335, 75–79.

Hughes, M.E., Grant, G.R., Paquin, C., Qian, J. and Nitabach, M.N. (2012) Deep sequencing the circadian and diurnal transcriptome of Drosophila brain. Genome Res., 22, 1266–1281.

Hughes, M.E., Hogenesch, J.B. and Kornacker, K. (2010) JTK-CYCLE: An efficient nonparametric algorithm for detecting rhythmic components in genome-scale data sets. J. Biol. Rhythms, 25, 372–380.

Ichimura, T. (1971) Sexual cell division and conjugation-papilla formation in sexual reproduction of Chlosterium strigosum. 7th Int. Seaweed Symp., 208–214. at: https://ci.nii.ac.jp/naid/10027352494/en/.

Kamioka, M., Takao, S., Suzuki, T., Taki, K., Higashiyama, T., Kinoshita, T. and Nakamichi, N. (2016) Direct Repression of Evening Genes by CIRCADIAN CLOCK-ASSOCIATED1 in the Arabidopsis Circadian Clock. Plant Cell, 28, 696–711. Available at: http://www.plantcell.org/lookup/doi/10.1105/tpc.15.00737.

Kester, D.R., Duedall, I.W., Connors, D.N. and Pytkowicz, R.M. (1967) Preparation of Artificial Seawater. Limnol. Oceanogr., 12, 176–179. Available at: https://aslopubs.onlinelibrary.wiley.com/doi/abs/10.4319/lo.1967.12.1.0176.

Lagercrantz, U. (2009) At the end of the day: A common molecular mechanism for photoperiod responses in plants? J. Exp. Bot., 60, 2501–2515.

Levin, M., Anavy, L., Cole, A.G., et al. (2016) The mid-developmental transition and the evolution of animal body plans. Nature, 531, 637–641.

Lien, T. and Knutsen, G. (1979) SYNCHRONOUS GROWTH OF CHLAMYDOMONAS REINHARDTII (CHLOROPHYCEAE): A REVIEW OF OPTIMAL CONDITIONS. J. Phycol., 15, 191–200.

Linde, A.-M., Eklund, D.M., Kubota, A., et al. (2017) Early evolution of the land plant circadian clock. New Phytol., 216, 576–590. Available at: http://www.ncbi.nlm.nih.gov/pubmed/28244104%0Ahttp://www.pubmedcentral.nih.gov/articlerender.fcgi?artid=PMC5638080.

Lohse, M., Nagel, A., Herter, T., et al. (2014) Mercator: A fast and simple web server for genome scale functional annotation of plant sequence data. Plant, Cell Environ., 37, 1250–1258.

los Reyes, P. de, Romero-Campero, F.J., Ruiz, M.T., Romero, J.M. and Valverde, F. (2017) Evolution of Daily Gene Co-expression Patterns from Algae to Plants. Front. Plant Sci., 8. Available at: http://journal.frontiersin.org/article/10.3389/fpls.2017.01217/full.

Michael, T.P., Breton, G., Hazen, S.P., Priest, H., Mockler, T.C., Kay, S.A. and Chory, J. (2008) A morning-specific phytohormone gene expression program underlying rhythmic plant growth. PLoS Biol., 6, 1887–1898.

Mittag, M., Kiaulehn, S. and Johnson, C.H. (2005) The circadian clock in Chlamydomonas reinhardtii. What is it for? What is it similar to? Plant Physiol., 137, 399–409.

Miyazaki, Y., Takase, T. and Kiyosue, T. (2015) ZEITLUPE positively regulates hypocotyl elongation at warm temperature under light in Arabidopsis thaliana. Plant Signal. Behav., 10, 1–3.

Mockler, T.C., Michael, T.P., Priest, H.D., Shen, R., Sullivan, C.M., Givan, S.A., Mcentee, C., Kay, S.A. and Chory, J. (2007) The diurnal project: Diurnal and circadian expression profiling, model-based pattern matching, and promoter analysis. In Cold Spring Harbor Symposia on Quantitative Biology. pp. 353–363.

Monnier, A., Liverani, S., Bouvet, R., Jesson, B., Smith, J.Q., Mosser, J., Corellou, F. and Bouget, F.Y. (2010) Orchestrated transcription of biological processes in the marine picoeukaryote Ostreococcus exposed to light/dark cycles. BMC Genomics, 11.

Mure, L.S., Le, H.D., Benegiamo, G., et al. (2018) Diurnal transcriptome atlas of a primate across major neural and peripheral tissues. Science (80-. )., 359.

Nakamichi, N., Kiba, T., Henriques, R., Mizuno, T., Chua, N.H. and Sakakibara, H. (2010) PSEUDO-RESPONSE REGULATORS 9, 7, and 5 Are Transcriptional Repressors in the Arabidopsis Circadian Clock. Plant Cell, 22, 594–605. Available at: http://www.plantcell.org/cgi/doi/10.1105/tpc.109.072892.

Nakayama, H., Sakamoto, T., Okegawa, Y., et al. (2018) Comparative transcriptomics with self-organizing map reveals cryptic photosynthetic differences between two accessions of North American Lake cress. Sci. Rep., 8.

Nohales, M.A. and Kay, S.A. (2016) Molecular mechanisms at the core of the plant circadian oscillator. Nat. Struct. Mol. Biol., 23, 1061–1069.

Nusinow, D.A., Helfer, A., Hamilton, E.E., King, J.J., Imaizumi, T., Schultz, T.F., Farré, E.M. and Kay, S.A. (2011) The ELF4-ELF3-"LUX complex links the circadian clock to diurnal control of hypocotyl growth. Nature, 475, 398–404.

Nystedt, B., Street, N.R., Wetterbom, A., et al. (2013) The Norway spruce genome sequence and conifer genome evolution. Nature, 497, 579–84. Available at: http://www.ncbi.nlm.nih.gov/pubmed/23698360.

Oberschmidt, O., Hucking, C. and Piechulla, B. (1995) Diurnal Lhc Gene-Expression Is Present in Many but Not All Species of the Plant Kingdom. Plant Mol. Biol., 27, 147–153.

Price, D.C., Chan, C.X., Yoon, H.S., et al. (2012) Cyanophora paradoxa genome elucidates origin of photosynthesis in algae and plants. Science (80-. )., 335, 843–847. Available at: http://www.ncbi.nlm.nih.gov/pubmed/22344442.

Proost, S., Bel, M. Van, Sterck, L., Billiau, K., Parys, T. Van, Peer, Y. Van de and Vandepoele, K. (2009) PLAZA: a comparative genomics resource to study gene and genome evolution in plants. Plant Cell, 21, 3718–3731.

Proost, S., Krawczyk, A. and Mutwil, M. (2017) LSTrAP: Efficiently combining RNA sequencing data into co-expression networks. BMC Bioinformatics, 18.

Rawat, R., Schwartz, J., Jones, M.A., Sairanen, I., Cheng, Y., Andersson, C.R., Zhao, Y., Ljung, K. and Harmer, S.L. (2009) REVEILLE1, a Myb-like transcription factor, integrates the circadian clock and auxin pathways. Proc. Natl. Acad. Sci., 106, 16883–16888. Available at: http://www.pnas.org/cgi/doi/10.1073/pnas.0813035106.

Rawat, R., Takahashi, N., Hsu, P.Y., Jones, M.A., Schwartz, J., Salemi, M.R., Phinney, B.S. and Harmer, S.L. (2011) REVEILLE8 and PSEUDO-REPONSE REGULATOR5 form a negative feedback loop within the arabidopsis circadian clock. PLoS Genet., 7.

Rensing, S.A., Lang, D., Zimmer, A.D., et al. (2008) The Physcomitrella genome reveals evolutionary insights into the conquest of land by plants. Science, 319, 64–9. Available at: http://www.sciencemag.org/content/319/5859/64.

Ritchie, M.E., Phipson, B., Wu, D., Hu, Y., Law, C.W., Shi, W. and Smyth, G.K. (2015) limma powers differential expression analyses for RNA-sequencing and microarray studies. Nucleic Acids Res., 43, e47.

Ronquist, F., Teslenko, M., Mar, P. van der, et al. (2012) MrBayes 3.2: efficient Bayesian phylogenetic inference and model choice across a large model space. Syst. Biol., 61, 539–542. Available at: http://sysbio.oxfordjournals.org/content/61/3/539.short.

Ruprecht, C., Mendrinna, A., Tohge, T., Sampathkumar, A., Klie, S., Fernie, A.R., Nikoloski, Z., Persson, S. and Mutwil, M. (2016) FamNet: A Framework to Identify Multiplied Modules Driving Pathway Expansion in Plants. Plant Physiol., 170, 1878–94.

Ruprecht, C., Proost, S., Hernandez-Coronado, M., Ortiz-Ramirez, C., Lang, D., Rensing, S.A., Becker, J.D., Vandepoele, K. and Mutwil, M. (2017) Phylogenomic analysis of gene co-expression networks reveals the evolution of functional modules. Plant J., 90.

Ruprecht, C., Vaid, N., Proost, S., Persson, S. and Mutwil, M. (2017) Beyond Genomics: Studying Evolution with Gene Coexpression Networks. Trends Plant Sci. Available at: http://www.ncbi.nlm.nih.gov/pubmed/28126286 [Accessed February 17, 2017].

Schaffer, R., Ramsay, N., Samach, A., Corden, S., Putterill, J., Carré, I.A. and Coupland, G. (1998) The late elongated hypocotyl mutation of Arabidopsis disrupts circadian rhythms and the photoperiodic control of flowering. Cell, 93, 1219–1229.

Serrano-Bueno, G., Romero-Campero, F.J., Lucas-Reina, E., Romero, J.M. and Valverde, F. (2017) Evolution of photoperiod sensing in plants and algae. Curr. Opin. Plant Biol., 37, 10–17.

Singh, R.K., Svystun, T., AlDahmash, B., Jï¿½nsson, A.M. and Bhalerao, R.P. (2017) Photoperiod-and temperature-mediated control of phenology in trees – a molecular perspective. New Phytol., 213, 511–524.

Stockel, J., Welsh, E.A., Liberton, M., Kunnvakkam, R., Aurora, R. and Pakrasi, H.B. (2008) Global transcriptomic analysis of Cyanothece 51142 reveals robust diurnal oscillation of central metabolic processes. Proc. Natl. Acad. Sci., 105, 6156–6161. Available at: http://www.pnas.org/cgi/doi/10.1073/pnas.0711068105.

Stöver, B.C. and Müller, K.F. (2010) TreeGraph 2: Combining and visualizing evidence from different phylogenetic analyses. BMC Bioinformatics, 11.

Suzuki, K., Ehara, T., Osafune, T., Kuroiwa, H., Kawano, S. and Kuroiwa, T. (1994) Behavior of mitochondria, chloroplasts and their nuclei during the mitotic cycle in the ultramicroalga Cyanidioschyzon merolae. Eur. J. Cell Biol., 63, 280–8. Available at: http://www.ncbi.nlm.nih.gov/pubmed/8082652.

Thain, S.C., Hall, a and Millar, a J. (2000) Functional independence of circadian clocks that regulate plant gene expression. Curr. Biol., 10, 951–6. Available at: http://www.ncbi.nlm.nih.gov/pubmed/10985381.

Thimm, O., Bläsing, O., Gibon, Y., et al. (2004) MAPMAN: A user-driven tool to display genomics data sets onto diagrams of metabolic pathways and other biological processes. Plant J., 37, 914–939.

Usadel, B., Bläsing, O.E., Gibon, Y., et al. (2008) Multilevel genomic analysis of the response of transcripts, enzyme activities and metabolites in Arabidopsis rosettes to a progressive decrease of temperature in the non-freezing range. Plant, Cell Environ., 31, 518–547.

Usadel, B., Blasing, O.E., Gibon, Y., Retzlaff, K., Hohne, M., Gunther, M. and Stitt, M. (2008) Global Transcript Levels Respond to Small Changes of the Carbon Status during Progressive Exhaustion of Carbohydrates in Arabidopsis Rosettes. PLANT Physiol., 146, 1834–1861. Available at: http://www.plantphysiol.org/cgi/doi/10.1104/pp.107.115592.

Vaulot, D., Marie, D., Olson, R.J. and Chisholm, S.W. (1995) Growth of Prochlorococcus, a photosynthetic prokaryote, in the equatorial Pacific Ocean. Science (80-. )., 268, 1480–1482.

Vazquez, A., Flammini, A., Maritan, A. and Vespignani, A. (2003) Global protein function prediction from protein-protein interaction networks. Nat. Biotechnol., 21, 697–700. Available at: http://www.nature.com/doifinder/10.1038/nbt825.

Wang, Z.Y. (1997) A Myb-Related Transcription Factor Is Involved in the Phytochrome Regulation of an Arabidopsis Lhcb Gene. PLANT CELL ONLINE, 9, 491–507. Available at: http://www.plantcell.org/cgi/doi/10.1105/tpc.9.4.491.

Wu, L.F., Hughes, T.R., Davierwala, A.P., Robinson, M.D., Stoughton, R. and Altschuler, S.J. (2002) Large-scale prediction of Saccharomyces cerevisiae gene function using overlapping transcriptional clusters. Nat. Genet., 31, 255–265. Available at: http://www.nature.com/doifinder/10.1038/ng906.

Xu, W., Yang, R., Li, M., et al. (2011) Transcriptome phase distribution analysis reveals diurnal regulated biological processes and key pathways in rice flag leaves and seedling leaves. PLoS One, 6.

Yakir, E., Hassidim, M., Melamed-Book, N., Hilman, D., Kron, I. and Green, R.M. (2011) Cell autonomous and cell-type specific circadian rhythms in Arabidopsis. Plant J., 68, 520–531.

Zhang, R., Lahens, N.F., Ballance, H.I., Hughes, M.E. and Hogenesch, J.B. (2014) A circadian gene expression atlas in mammals: implications for biology and medicine. Proc Natl Acad Sci U S A, 111, 16219–16224.

Zhou, L., Cheng, D., Wang, L., Gao, J., Zhao, Q., Wei, W. and Sun, Y. (2017) Comparative transcriptomic analysis reveals phenol tolerance mechanism of evolved Chlorella strain. Bioresour. Technol., 227, 266–272.

Zones, J.M., Blaby, I.K., Merchant, S.S. and Umen, J.G. (2015) High-Resolution Profiling of a Synchronized Diurnal Transcriptome from Chlamydomonas reinhardtii Reveals Continuous Cell and Metabolic Differentiation, Available at: http://www.plantcell.org/content/early/2015/10/02/tpc.15.00498.abstract?papetoc.

